# Conjunctive and complementary CA1 hippocampal cell populations relate sensory events to immobility and locomotion

**DOI:** 10.1101/2022.07.06.498996

**Authors:** Samsoon Inayat, Brendan B. McAllister, Ian Q. Whishaw, Majid H. Mohajerani

## Abstract

This study investigated the dynamics of recruitment of cells in the CA1 region of the hippocampus in response to sensory stimuli presented during immobility, movement, and their transitions. Two-photon calcium imaging of somal activity in CA1 neuron populations was done in head fixed mice. Sensory stimuli, either a light flash or an air stream, were delivered to the mice when at rest, when moving spontaneously, and while they were induced to run a fixed distance on the conveyor belt. Overall, 99% of 2083 identified cells (from 5 mice) were active across one or more of 20 sensorimotor events. A larger proportion of cells were active during locomotion. Nevertheless, for any given sensorimotor event, only about 17% of cells were active. When considering pairs of sensorimotor event types, the active cell population consisted of conjunctive (C ∈ A and B) cells, active across both events, and complementary (C ∈ A not B or C ∈ B not A) cells that were active only during individual events. Whereas conjunctive cells characterised stable representations of repeated sensorimotor events, complementary cells characterised recruitment of new cells for encoding novel sensorimotor events. The moment-to-moment recruitment of conjunctive and complementary cells across changing sensorimotor events signifies the involvement of the hippocampus in functional networks integrating sensory information with ongoing movement. This role of the hippocampus is well suited for movement guidance that secondarily might include spatial behavior, episodic learning and memory, context representation, and scene construction.

## Introduction

The idea that the brain features behavior-specific electrographic events during the behavioral states of immobility and locomotion originated with the finding that electrical activity in the hippocampus is different during these two states (Kay and Frank, 2019; Vanderwolf, 1969). The electrical activity measured as local field potentials (LFPs) features large amplitude irregular activity (LIA) during immobility and two kinds of rhythmical slow activity (RSA or theta activity, 6-12 Hz) that are sensory or movement related (Buzsáki, 2006; Buzsaki and Watson, 2012; Foster et al., 1989; Perentos et al., 2022; Vanderwolf, 1969; Whishaw, 1976; Whishaw and Vanderwolf, 1971, 1973). Sensory-induced RSA is cholinergic-dependent whereas movement-related RSA can be cholinergic-independent (Kramis et al., 1975; Vanderwolf et al., 1984; Whishaw and Dyck, 1984). The behavior-specific differentiation of electrical activity is also found at the cellular level. Whereas excitatory cells in both CA1 and CA2 modulate their firing in relation to location during immobility (Kay et al., 2016; Yu et al., 2017) and during locomotion (O’Keefe and Dostrovsky, 1971; O’Keefe and Nadel, 1978), anatomically distinct types of inhibitory cells fire preferentially during locomotion versus immobility (Arriaga and Han, 2017). Location specific firing during locomotion has also been found in other subregions of the hippocampal formation such as the dentate gyrus, CA3, subiculum, and medial entorhinal cortex, as well as cortical regions including the retrosplenial, visual, and sensorimotor cortices. There are reports of behavior-dependent cellular responses in the hippocampus when animals encounter sensory stimuli including odors (Igarashi et al., 2014; Komorowski et al., 2009; MacDonald et al., 2013; Manns et al., 2007; Taxidis et al., 2020), tones (Moita et al., 2003; Sakurai, 2002), tastes (Herzog et al., 2019), textures (Itskov et al., 2011), visual stimuli (Chen et al., 2013a; Zhao et al., 2020), somatosensory stimuli (Bellistri et al., 2013; Wang et al., 2014), and vestibular events (Smith, 1997).

Behavior-specific sensory processing is also reported in sensory and non-sensory cortices, and some subcortical structures display behavior-dependent spontaneous and evoked firing activity - for a review, see (Schneider, 2020). Compared to immobility, locomotion increases firing of sensory neurons in the visual (Dipoppa et al., 2018; Niell and Stryker, 2010) and somatosensory (Ayaz et al., 2019; Sofroniew et al., 2015) cortices but reduces firing in the auditory cortex (Schneider et al., 2014). Locomotion also modulates higher order functions; for example, it improves sensory perception (Mineault et al., 2016), guides navigation through the visual cortex (Saleem et al., 2018), and alters the functional connectivity of the visual and retrosplenial cortices with other cortical regions (Clancy et al., 2019). Movements other than locomotion, such as whisking, facial and body movements, and arousal-related changes in pupil diameter, also influence neuronal firing in the sensory cortices (Musall et al., 2019; Stringer et al., 2019). Furthermore, movement-related inputs to the sensory cortices reflect expectations of sensory information anticipated by movements (Attinger et al., 2017; Keller et al., 2012). These studies have contributed to the idea that such activity mediates spatial navigation and context processing [see reviews, (Colgin, 2020; Eichenbaum et al., 2016; Moser et al., 2017; Smith and Bulkin, 2014)].

Stimulated by the many findings of the relationship between electrophysiological events and ongoing behavior, the present study examined behavior-specific hippocampal cellular responses to sensory stimuli, specifically focusing on how cellular responses are modulated during immobility, locomotion, and their transitions. Calcium imaging was used in head-fixed mice to record hippocampal CA1 cell responses to sensory stimulation by a pressurized air stream or light flash while the mice were at rest or locomoting. A population level analysis investigated how cells were recruited during and between sensory stimuli applied during immobility versus locomotion. Subgroups of cells responding to different sensorimotor events were active in different stimulus-locomotion configurations. When two (or more) events were considered at a time, cells could be classified as complementary, responsive to only one event, or conjunctive, responsive to both events. From a large pool of responsive cells, complementary cells identified novel sensorimotor events, and conjunctive cells encoded familiar sensorimotor events, suggesting a recruitment with replacement organization.

## Results

### Experimental design, behavioral paradigm, and its characterization

Hippocampal CA1 cell activity was recorded from head-fixed mice resting or moving on a non-motorized conveyor belt (Fig. 1A). The application of a brake restricted movement, and its release enabled movement and running. An air stimulus, delivered on the back, and whole-field illumination with an LED in front of the mouse served as sensory stimuli. The behavioral paradigm featured 7 configurations of motor activity and sensory stimulation (Fig. 1B), divided into two sets based on two brake conditions: Brake (B) and No-Brake (NB). In Configurations 1, 2, 6, and 7, the B-condition prevented walk/run behaviors, whereas in Configurations 3, 4, and 5, the NB-condition allowed walk/run behaviors. In Configurations 1 and 2, ten pulses each of light (L; 200 ms with 5 s interval) and air (A; 5 s with 10 s interval) were applied, respectively. In Configurations 3, 4, and 5, the air stream was applied to cause stimulus-induced running for a fixed distance of 150 cm (i.e., one lap of the belt), after which the air was stopped. Ten trials were given in each Configuration, with an intertrial interval of 15 s. In Configuration 4, a 200 ms light flash was also applied once the mouse had traveled 110 cm. Configurations 3 and 5, 1 and 6, and 2 and 7 were identical.

**Figure 1.**
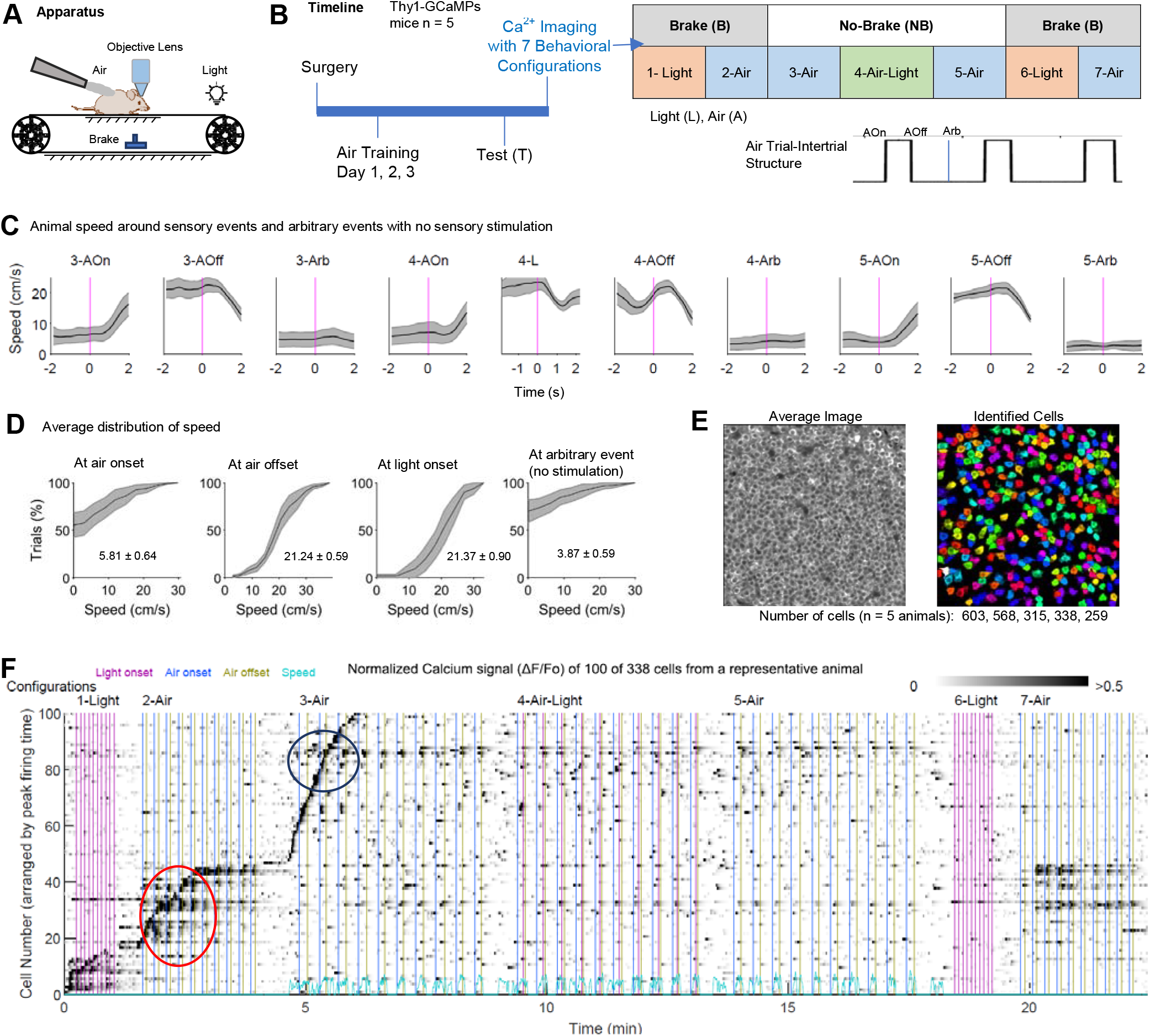
The experimental paradigm for studying hippocampal CA1 neuronal responses to sensory stimulation during immobility and locomotion. **(A)** Experimental setup. A head-fixed mouse on a linear conveyor belt receives a mild air stream on the back to motivate running. **(B)** Experiment timeline and list of 7 configurations that mice experienced while calcium imaging was performed. Inset shows schematic depicting air onset (AOn), air offset (AOff), and arbitrary (Arb) events. **(C)** Average speed over all trials by animals around stimulus and arbitrary (Arb) events during Configurations 3, 4, and 5. Magenta lines indicate occurrences of events. **(D)** Distribution of speeds at events mentioned in C. The numbers indicate overall mean±SEM speeds. **(E)** Average image (left) of the calcium imaging time series from a representative animal, and arbitrarily color-coded regions of interest indicating identified cell bodies (right) in a ~400 × 400 μm imaging window. **(F)** Normalized calcium traces (ΔF/F_0_), from 100 representative cells from a representative animal for a whole recording of ~25 min, overlayed with indicators of light and air stimuli (vertical-colored lines). Cells are arranged by the time of peak firing. The speed signal is shown at the bottom (cyan color). Notice differential firing of cells during immobility vs. locomotion (cells in blue vs red ovals) and cell population changes from sensorimotor event to sensorimotor event. Thick black lines in (C) and (D) indicate mean over animals while shaded regions represent SEM.

Neuronal activity 2 s before and 2 s after each stimulus event was analyzed and compared across B and NB conditions. For air stimuli, activity around both air onset and offset events (AOn and AOff respectively) was characterized, whereas for light stimuli activity was analyzed only at light onset (L) because of the brevity of the stimulus duration (i.e., 200 ms). Because the hippocampus is an associative brain region, cellular activity may have resulted from intrinsic network activity representing task parameters or internal states. Therefore, the activity of cells was also analyzed around arbitrary (Arb) time points at which no explicit sensory stimulation occurred for both the B and NB conditions. The time chosen for the Arb events (Fig. 1B, inset) was the mid-point of the intertrial intervals (in Configurations 2, 3, 4, 5, and 7). For the B and NB conditions, the Arb point was 5 s and 7.5 s after air offset, respectively. Finally, during the NB-condition, activity of neurons was analyzed when animals voluntarily started or stopped their locomotion in the absence of explicit sensory stimuli.

The animals were immobile during the B-condition. During the NB-condition, their average speed was always greater than zero around stimulus and Arb events (Fig. 1C). For AOff and L events, the speed of individual animals in all trials (from Configurations 3-5) was greater than zero (Fig. 1D, middle panels). For AOn, however, speeds were greater than zero on 50.00 ± 12.78 % of the trials (average over 5 animals) had speeds greater than zero (Fig. 1D, left panel). There was animal-to-animal variability, with greater than zero speeds in 3, 26, 18, 17, and 11 trials (corresponding to 5 animals) of the cumulative 30 air application trials. In trials in which the animals were immobile at AOn, the animals initiated locomotion after air onset with an average movement latency of 0.94 ± 0.06 sec (range: 0.05, 3.66, median: 0.62). Hence, for subsequent analyses, these trials were pooled with trials in which the mice were already moving at air onset. For the Arb events, there was also animal-to-animal variability, with greater than zero speeds in 2, 20, 21, 8, and 6 trials (corresponding to 5 animals) of the cumulative 30 trials (Fig. 1D). Overall, however, the average speed around the Arb events was greater than zero (Fig. 1C).

### Calcium imaging of the activity of hippocampal CA1 pyramidal neurons

Calcium imaging of CA1 pyramidal neurons was performed as the mice experienced the 7 behavioral configurations. Fig. 1E presents an average image of an imaging stack from a representative animal, showing densely packed neurons in dorsal CA1. Using Suite2p, a Python based analysis software, 603, 568, 315, 338, and 259 individual cells (total: 2083 cells) were identified from 5 mice. An image of identified cells from a representative animal is displayed in Fig. 1E. The figure also shows that Suite2P did not recognize all visible cells in the average image, perhaps because some cells were seldom active (fired only occasionally or not at all). The raw calcium traces (ΔF/F_0_) of 100 cells from a representative animal shown in Fig. 1F demonstrate that cellular responses reflected brake conditions and/or sensorimotor events. For one example, the cell groups colored red were active in the B-condition, and those colored blue were active in the NB-condition. For further quantitative analysis of cellular responses to sensory events and behavior, as well as study of population level characteristics, the calcium signals were deconvolved to estimate firing rates for each cell. In the subsequent text, neural activity refers to the firing rates estimated from the calcium signals.

To examine the changes in cellular firing properties across the B and NB conditions, mean and maximum firing rates were determined for both conditions and for each cell. The mean and maximum firing rates were then calculated for each animal by averaging the respective values across cells. These values were then subjected to statistical comparison across conditions with a repeated measures analysis of variance (RM-ANOVA). The mean firing rate across the B and NB conditions was not significantly different [0.054 ± 0.016 vs. 0.073 ± 0.022; *F* (1,4) = 7.05, *p* = .057, η^2^ = .64], but note a higher mean for the NB-condition and a *p*-value close to .05. The comparison of the maximum firing rates of the cells for the B and NB conditions gave a significant difference [26.47 ± 5.05 vs. 48.51 ± 10.14; *F* (1,4) = 13.89, *p* = .020, η^2^ = .78] showing that locomotion contributed to a transient increase in cell firing.

### Larger population of responsive cells for No-Brake condition versus Brake condition for AOn, AOff, and Arb events

Cell activity during the B and NB conditions for AOn, AOff, and Arb events was investigated by finding responsive cells and their response characteristics. Cells active around these events were examined by generating raster plots of neuronal activity (trials versus time) for each cell and event. Peri-event time histograms (PETHs) were then calculated from the raster plots by finding mean neuronal activity over trials. Responsive cells were identified from PETHs by statistically comparing average pre- and post-event firing rates using a Student’s t-test (α = .05). This method allowed an unbiased approach for identifying both excited (Exc) and inhibited (Inh) cells, in which post-event firing rates were significantly larger or smaller, respectively, compared to pre-event firing rates. Figure 2A displays raster plots for a representative AOn-Exc cell (left) and AOn-Inh cell (right) from Configuration 2, with firing rates across time and across trials overlayed with the PETH. Population level response dynamics for responsive cells were observed with rate vector maps (Fig. 2B, top row). These show the normalized mean response of all cells organized by the time of each cell’s peak response. The rate vector maps for both the B and NB conditions highlight the binary nature of Exc and Inh cells (i.e., their responses of ON or OFF type) in relation to AOn. This can also be observed in the population correlation plots (Fig. 2B, middle row), in which each pixel represents a Pearson correlation between two columns of the corresponding rate vector map. The population correlation plots displayed dark square-like regions for both the pre-event (−2 to 0 s) and post-event (0 to 2 s) periods, corresponding to responses of Exc and Inh cells, respectively. Plots of the average population correlation (Fig. 2B, bottom row) confirmed that the majority of Exc and Inh cells were binary in nature. Plots generated for AOff and Arb events exhibited similar characteristics, with both Exc and Inh cells.

**Figure 2.**
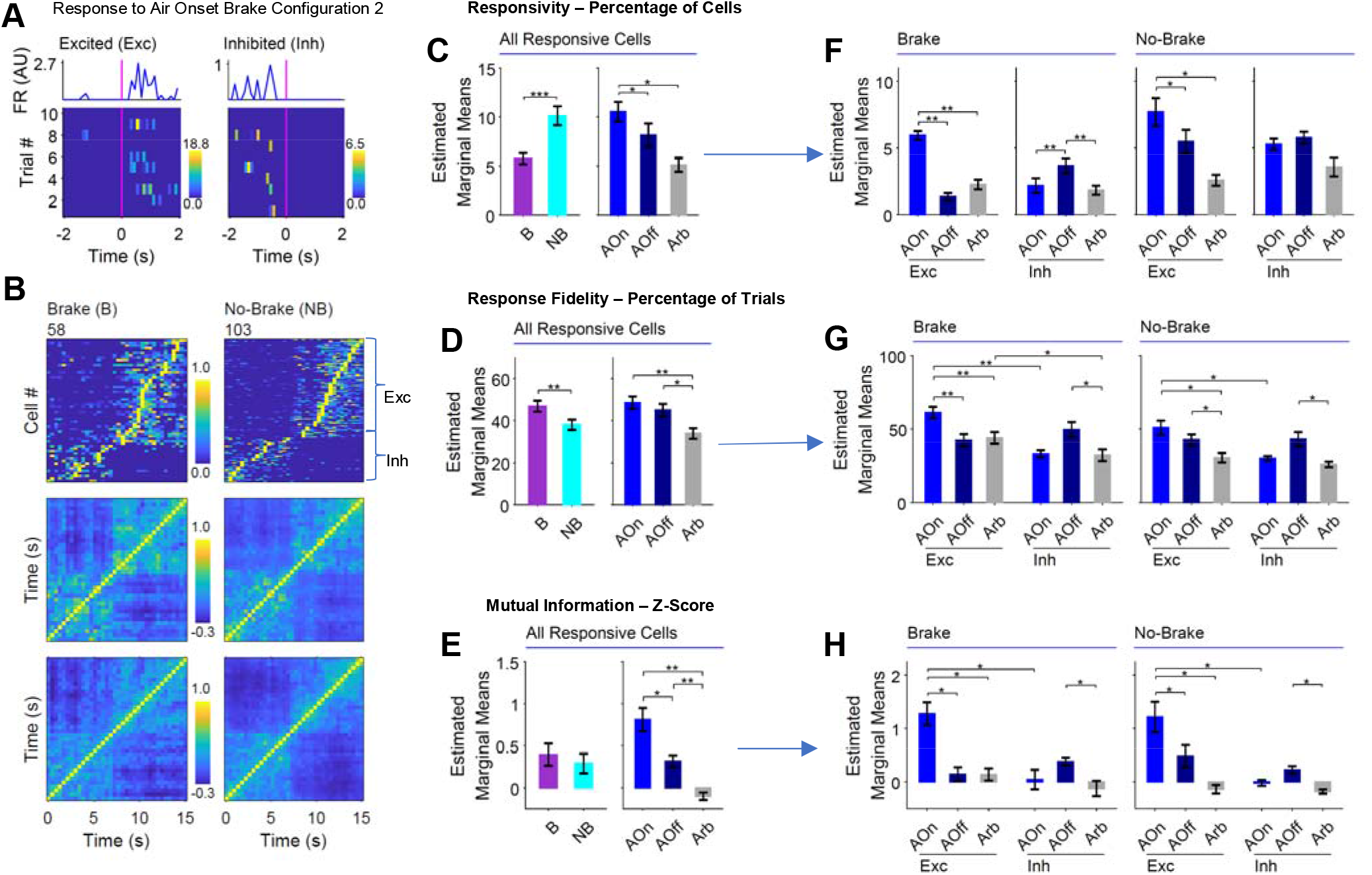
More cells active during the No-Brake compared to the Brake condition for AOn, AOff, and Arb events. **(A)** Raster plots and peri-event histograms of representative excited (Exc) and inhibited (Inh) cells showing increased and decreased firing, respectively, in response to AOn (t = 0, magenta line). The blue lines show the means across trials. **(B)** Rate vector maps (top row) and corresponding population vector correlation plots (middle row) of responsive cells to AOn from a representative animal for Brake vs. No-Brake conditions. Average population correlation plots (bottom row) from 5 animals. **(C-E)** Results of two-way RM-ANOVA for assessing responsivity (Resp), response fidelity (RF), and mutual information z-score (zMI) respectively, across brake conditions (B versus NB) and events (Aon, AOff, and Arb) **(F-H)** Results of three-way RM-ANOVA for assessing Resp, RF, and zMI respectively, across brake conditions, events, and cell types (Exc vs Inh). For all statistical tests, *n* = 5 mice. Error bars = SEM. \**p* < .05, \*\**p* < .01, \*\*\**p* < .001.

To assess the responsivity (Resp) of cell activation across the B and NB conditions, the percentage of cells defined as responsive (including both Exc and Inh cells) was found for the AOn, AOff, and Arb events. Resp was averaged over Configurations 2 and 7 and over Configurations 3, 4, and 5 for the B and NB conditions, respectively. The data were subjected to a RM-ANOVA with two within-subjects factors (Brake-State and Event-Type). There were significant main effects of both Brake-State [*F* (1,4) = 130.12, *p* < .001, η^2^ = .97] and Event-Type [*F* (2,8) = 19.22, *p* = .009, η^2^= .83] with no interaction. The Resp in the NB-condition was larger than in the B-condition, and *post hoc* comparisons revealed that the percentage of cells activated around AOn was larger than for the AOff and Arb events (Fig. 2C).

Responsive cells found from PETHs might have had different response characteristics across trials during B and NB conditions in response to AOn, AOff, and Arb events. Therefore, two additional measures, response fidelity (RF) and mutual information (MI), were extracted from the raster plots to quantitatively assess the strength and robustness of cellular responses across trials. RF was defined as the percentage of trials in which a cell’s firing rate was greater than zero at any point during a trial. MI between firing rate and time was determined from a raster plot (Souza et al., 2018) and then z-scored using a distribution of 500 MI values calculated from randomly shuffled instances of the same raster plot, giving a standardized measure of MI (zMI). A larger zMI value indicates greater temporal consistency of responses across trials. RF and zMI values were averaged across Configurations 2 and 7 and across Configurations 3, 4, and 5 for the B and NB conditions, respectively. The results were subjected to a RM-ANOVA with two within-subjects factors (Brake-State and Event-Type) individually for RF and zMI. For RF, there were significant main effects of both Brake-State [*F* (1,4) = 41.32, *p* = .003, η^2^ = .91] and Event-Type [*F* (2,8) = 30.62, *p* = .000, η^2^ = .88] with no interaction. RF in the B-condition was greater compared to the NB-condition, and *post hoc* comparisons revealed that the RF for AOn and AOff was greater compared to the Arb events (Fig. 2D). For zMI, there was a main effect of Event-Type [*F* (2,8) = 42.50, *p* = .001, η^2^ = .91], with *post hoc* comparisons showing that zMI for the AOn and AOff events was larger than for the Arb events, and also that zMI for the AOn events was larger than for AOff (Fig. 2E).

Taken together, the results suggest that more cells were active during the NB-condition compared to the B-condition. Nevertheless, cells that were active during the B-condition had more robust responses than occurred for cells active during the NB-condition. Additionally, cells that were activated by sensory air events had a more robust response compared to those activated around Arb events, especially at AOn.

### Characteristics of Exc and Inh cells are similar across Brake and No-Brake conditions but different for AOn and AOff events

The analysis of cell populations activated across B and NB conditions around AOn, AOff, and Arb events was further extended to address whether Brake-State and Event-Type evoked varying responses in the different cell types (i.e., Exc and Inh cells). A RM-ANOVA with three within-subjects factors (Brake-State, Event-Type, and Cell-Type) was used on the output variables: Resp, RF, and zMI.

For Resp, there was a significant three-way interaction [*F* (2,8) = 6.73, *p* = .028, η^2^ = .63].

Therefore, simple two-way interactions of two factors at each level of the third factor were assessed, using a Bonferroni adjusted alpha value (see methods). Significant interactions were found between Event-Type and Cell-Type for both the Brake [*F* (2,8) = 51.37, *p* < .001, η^2^ = .93] and the NB-condition [*F* (2,8) = 9.14, *p* = .010, η^2^ = .70]. In the next level of analysis, simple simple main effects were assessed, again using a Bonferroni adjusted alpha value. Only effects of Event-Type were significant; in the B-condition there was a simple simple main effect for both the Exc cells [*F* (2,8) = 56.15, *p* < .001, η^2^ = .93] and the Inh cells [*F* (2,8) = 32.19, *p* < .001, η^2^ = .89], and in the NB-condition, there was a simple simple main effect for only the Exc cells [*F* (2,8) = 12.98, *p* = .003, η^2^ = .76]. Simple simple comparisons (Bonferroni-corrected) were then used to assess differences in means for the Event-Types (Fig. 2F). For the Exc cells, AOn events generated stronger responses (greater Resp) compared to AOff or Arb events in both the B and NB conditions. For the Inh cells, however, AOff events were the most potent at increasing Resp, specifically in the B condition, though a similar but non-significant trend was observed in the NB condition.

For RF, there was a main effect of Brake-State [*F* (1,4) = 32.21, *p* = .005, η^2^ = .89] and a twoway interaction between Event-Type and Cell-Type [*F* (2,8) = 25.50, *p* = .006, η^2^ = .86]. *Post hoc* comparisons (Fig. 2G) showed that, in both the B and NB conditions, Exc cells exhibited the greatest RF for AOn events, whereas Inh cells exhibited the greatest RF for AOff events. For zMI, there was a significant two-way interaction between Event-Type and Cell-Type [*F* (2,8) = 21.26, *p* = .004, η^2^ = .84]. *Post hoc* comparisons revealed that, similar to RF, zMI was greatest for the Exc cells around the AOn events and greatest for the Inh cells around the AOff events (Fig. 2H).

Taken together, the results suggest that the Exc and Inh cells were similar in the B and NB conditions, but AOn was associated with more Exc cell responses, and AOff was associated with more Inh cell responses. As found before, sensory air events generated more robust responses compared to Arb events for both the Exc and Inh cells.

### Separate populations of cells in Brake and No-Brake conditions for AOn, AOff, and Arb events

The population level activity of cells was classified across brake conditions and different events as conjunctive or complementary. Considering two sensorimotor events (e.g., A and B), conjunctive (Conj) cells were defined as those active for both events (Conj ∈ A and B), and complementary (Comp) cells were defined as those active for individual events. Comp cells could therefore be broken down into two populations (Comp1 ∈ A not B; and Comp2 ∈ B not A). The percentage of the total number of detected cells that was classified into each of three populations (Conj, Comp1, and Comp2) for different sensorimotor events. The percentages were then compared statistically or subjected to an agglomerative hierarchical clustering analysis to observe relations among sensorimotor events and configurations.

Comp1, Conj, and Comp2 cell populations were found for AOn, AOff, and Arb events, where Comp1 and Comp2 were selectively active during the B and NB conditions, respectively, and Conj were active in both. For this analysis, cells were pooled across Configurations 2, 7 and Configurations 3, 4, 5 for the B and NB conditions, respectively. The population percentage values were then subjected to a two-way RM-ANOVA, with Event-Type and Population-Type as within-subjects factors. There was a significant main effect of both Event-Type [*F* (2,8) = 10.53, *p* = .030, η^2^ = .72] and Population-Type [*F* (2,8) = 715.73, *p* < .001, η^2^ = .99] but no significant interaction. The effect of Event-Type was expected, based on the differences in Resp across sensory air and Arb events described above (Fig. 2C and Fig. 3A). The effect of Population-Type was more interesting, however, as it showed differences in the number of cells recruited into each of the three populations, with Conj cells forming a significantly smaller population compared to Comp1 and Comp2, and with Comp2 forming a significantly larger population compared to Comp1, as identified with *post hoc* comparisons (Fig. 3A).

**Figure 3.**
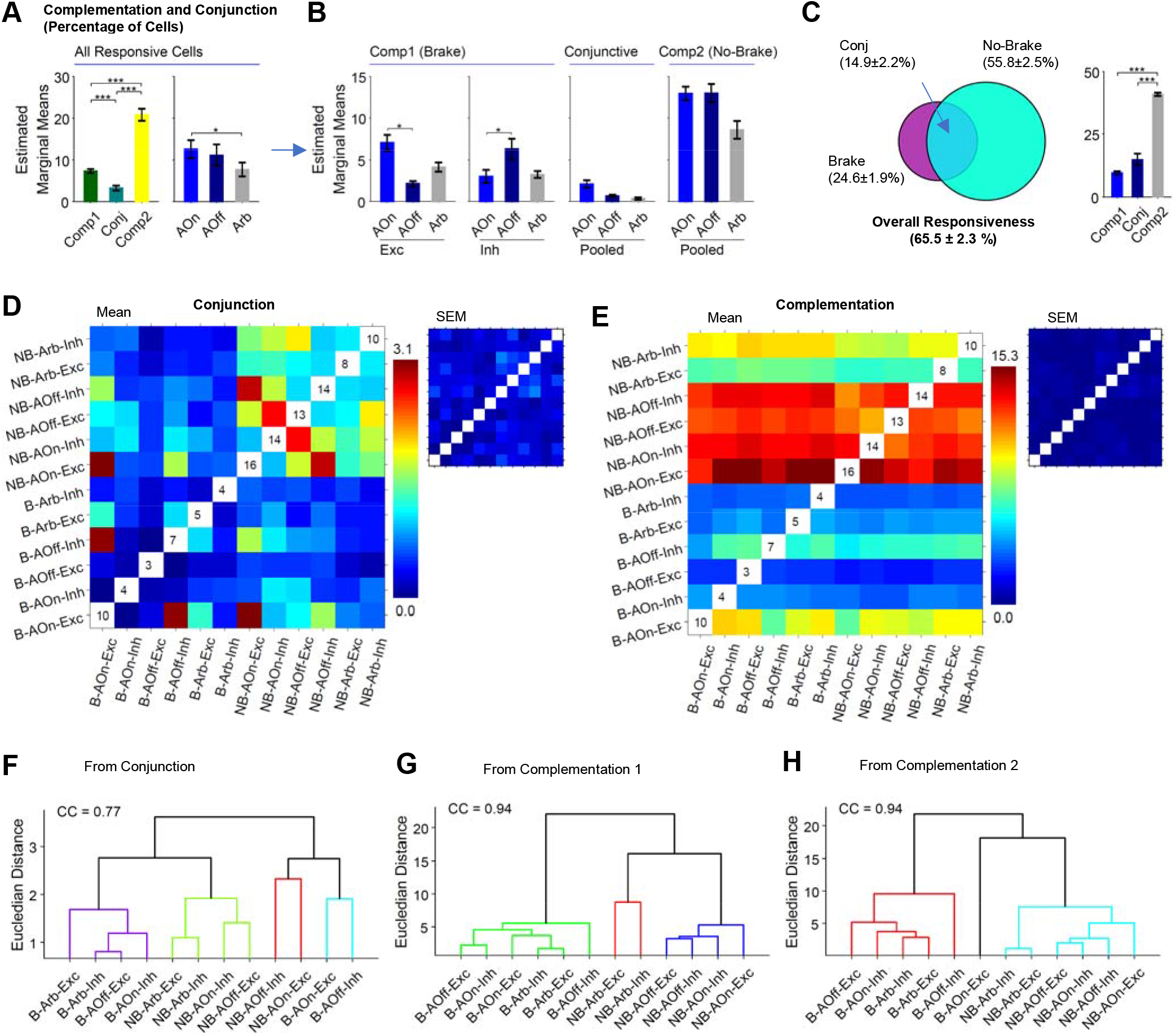
Segregated cell populations in the Brake and the No-Brake conditions for AOn, AOff, and Arb events. **(A)** Results of a two-way RM-ANOVA for assessing the percent of cells across population types (Comp1, Comp2, and conjunctive) and events (AOn, AOff, and Arb). **(B)** Results of a three-way RM-ANOVA assessing the percent of cells across cell types (Exc vs Inh) in addition to factors in (A). **(C)** Overall responsivity, conjunction, and complementation across Brake and No-Brake conditions. **(DE)** Average and SEM (insets) heatmaps (over 5 animals) of conjunction and complementation of Exc and Inh cell groups active in Brake and No-Brake conditions around AOn, AOff, and Arb events. **(F-H)** Dendrograms resulting from agglomerative hierarchical clustering of the average conjunction and complementation heatmaps/matrices. CC indicates cophenetic correlation. For all statistical tests, *n* = 5 mice. Error bars = SEM. \**p* < .05, \*\**p* < .01, \*\*\**p* < .001.

The complementation and conjunction of Exc and Inh cells was then examined by adding a third factor (Cell-Type) to the analysis and performing a three-way RM-ANOVA. There was a significant three-way interaction [*F* (4,16) = 7.19, *p* = .002, η^2^ = .64]. Further tests showed that the only simple two-way interaction was between Cell-Type and Event-Type for the Comp1 population [*F* (2,8) = 20.68, *p* = .003, η^2^ = .84]. Simple simple main effects of both Cell-Type and Event-Type were significant [*F* (2,8) = 12.88, *p* = .003, η^2^ = .76; and *F* (2,8) = 11.31, *p* = .005, η^2^ = .74, respectively]. *Post hoc* comparisons showed that, for Comp1 Exc cells, a larger percentage of cells was active around AOn compared to AOff. The opposite relation was observed for Comp1 Inh cells (Fig. 3B). For the Conj and Comp2 populations, the percentage of active Exc and Inh cells did not differ across event types (Fig. 3B, pooled data).

When pooled across event and cell types, 65.5 ± 2.3 % of the detected cells were responsive, comprising three populations. One population was active during the B-condition (Comp 1, 9.7 ± 0.7 %), a second population was active during the NB-condition (Comp 2, 40.9 ± 0.6 %) and the third population was active in both conditions (Conj, 14.9 ± 2.2 %) (see Fig. 3C, Venn diagram). After pooling, however, the average percent of Comp1 cells was not significantly different from the average percentage of Conj cells (Fig. 3C, RM-ANOVA and *post hoc* comparisons). This suggests that some cells might have shared responsiveness across sensory air and Arb events. To further investigate this possibility, the conjunction and complementation of all different groups of cells was analyzed. The conjunction and complementation of different groups of cells categorized by brake condition (B vs. NB), event type (AOn vs. AOff vs. Arb), and cell type (Exc vs. Inh) were plotted as heatmaps. Figures 3D and 3E show average (from 5 animals) conjunction and complementation heatmaps, respectively, and the insets show SEM heatmaps. In the average heatmaps, diagonal pixels are shown as white with no color coding. The numbers therein represent the percentage of responsive cells for the selected category. Off-Diagonal pixels in these heatmaps show percentages of conjunctive and complementary cells. For the complementation heatmap, pixels above and below the diagonal represent Comp1 and Comp2 cells, respectively, when considering row-column pairs.

Conjunctive cell populations were small. Notably, the highest degree of conjunction (~3 % conjunction) was observed between Exc cells responsive to AOn and Inh cells responsive to AOff during the Brake conditions (Fig. 3D, B-AOn-Exc vs B-AOff-Inh, row 1 column 4). To classify cell populations and their relationships with each other, a secondary analysis was conducted using agglomerative hierarchical clustering (see methods) of the average heatmap matrix, revealing four distinct clusters (Fig. 3F). The cyan cluster grouped Exc cells responding to AOn with Inh cells responding to AOff during the B-condition. In other words, this cluster included cells that responded during air ON periods (i.e., when the air stream was on). A similar relation existed during the NB-condition (red cluster). The third cluster (purple) was constituted by cells from the B-condition that were active during air OFF periods (i.e., when the air stream was not on). The fourth cluster (green) represented air OFF cells for the NB-condition. These results show that conjunctive cells across sensory air and Arb events could be segregated into two groups based on whether the brake was on or off, and further subdivided into cells that responded to air ON and OFF periods.

The complementation heatmap showed segregation between the B and NB conditions, with the greatest degree of complementation between the NB-AOn-Exc and B-AOff-Exc cell groups (~15.3 %). The complementation heatmap was split into two heatmaps, symmetric across the diagonal, by using the upper and lower triangular matrices corresponding to the Comp1 and Comp2 populations. The clustering of the Comp1 heatmap (Fig. 3G) showed that cells were grouped separately across the B (green cluster) and NB (blue and red clusters) conditions. The clustering of Comp2 (Fig. 3H) showed similar cluster segregation based on the B and NB conditions (cyan and red clusters, respectively). However, within the dendrograms of event-type/cell-type combinations, grouping mainly occurred based on event-type. For example, NB-AOff-Exc was closer to NB-AOff-Inh, and NB-AOn-Exc was closer to NB-AOn-Inh. Taken together, the results show segregation of cell groups across the B and NB conditions when cells were divided into subgroups based on AOn, AOff, and Arb events and Exc and Inh cell types.

### Separate populations of cells for Light-onset in the Brake and No-Brake conditions

Light-onset cell populations were analyzed to investigate whether the same or different cellular networks were involved in the B and NB conditions. The above-mentioned analysis (for AOn, AOff, and Arb events) was repeated for cell populations active around light stimulus events in Configurations 1 and 4 for the B and NB conditions, respectively. Both Exc and Inh cells were found in a similar way as is explained above. Fig. 4A shows raster plots and PETHs for representative Exc and Inh cells from Configuration 1. Rate vector maps (Fig. 4B, top row) and population vector correlation plots (Fig. 4B, middle and bottom rows) showed the binary nature of cellular responses for both representative populations in the B and NB conditions.

**Figure 4.**
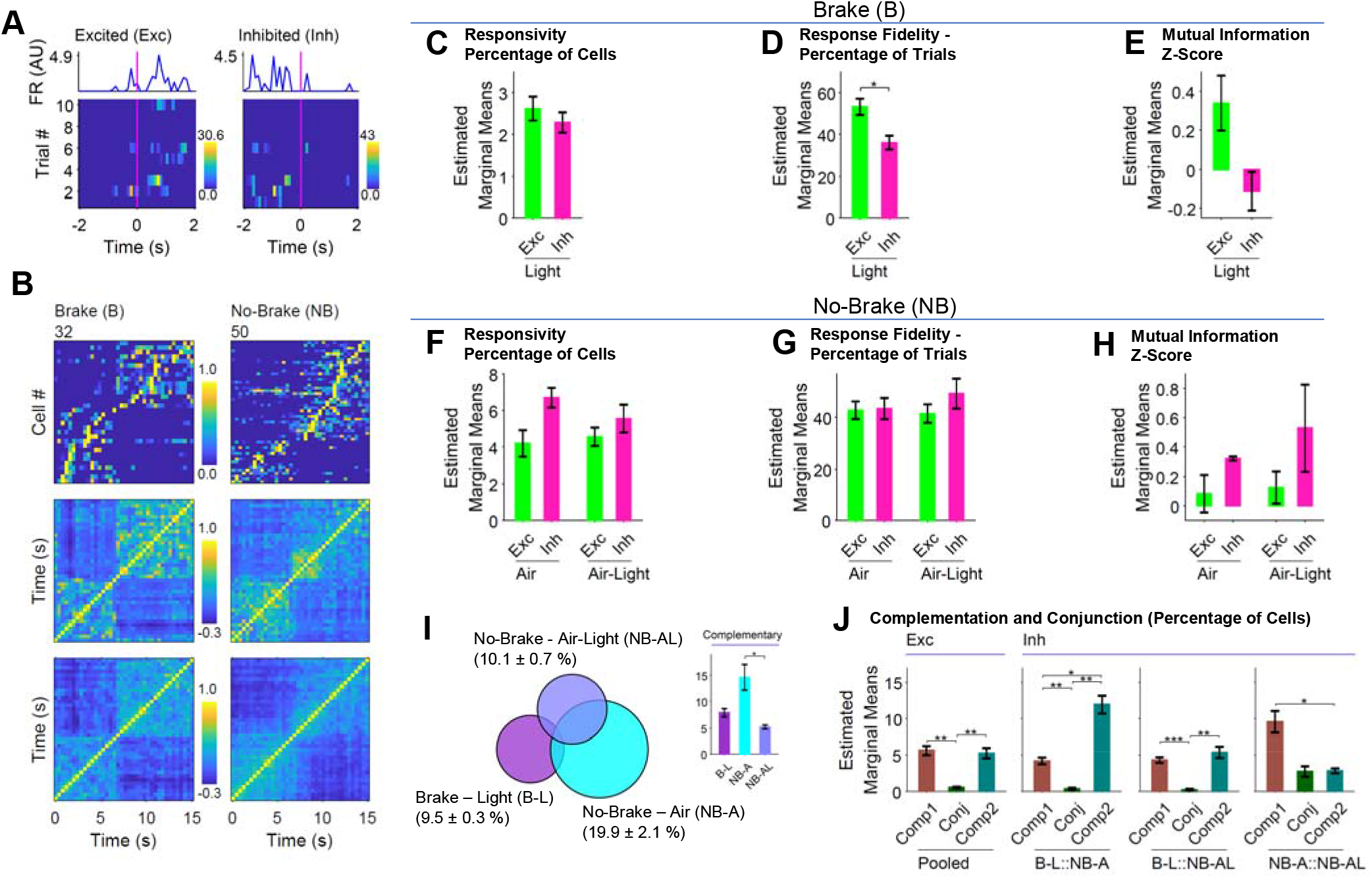
Segregated cell populations across the Brake and No-Brake conditions to a light stimulus. **(A)** Raster plots of representative Exc and Inh cells. The magenta line at t = 0 indicates light onset. The blue lines show means over trials. **(B)** Rate vector maps (top row) and corresponding population vector correlation plots (middle row) of responsive cells from a representative animal for Brake and No-Brake conditions. Average population correlation plots (bottom row) from 5 mice. **(C-E)** Results of one-way RM-ANOVA assessing responsiveness, response fidelity, and mutual information z-score of Exc and Inh cells for light stimulus in Brake condition (B-L). **(F-H)** Results of two-way RM-ANOVA assessing response variables for Exc and Inh cells in the No-Brake conditions for light stimulus combined with air (NB-AL) and identical Arb event where only air was present (NB-A). **(I)** Overall responsivity, conjunction, and complementation for B-L, NB-AL, and NB-A cell populations. **(J)** Results of threeway RM-ANOVA for assessing the percent of cells across population types (Comp1, Comp2, and Conj), cell types (Exc vs Inh) and event pairs e.g., B-L::NB-AL. For all statistical tests, *n* = 5 mice. Error bars = SEM. \**p* < .05, \*\**p* < .01, \*\*\**p* < .001.

Resp, RF, and zMI were separately assessed for the B and NB conditions, because the light stimulus was applied alone in the B-condition (B-L), whereas in the NB-condition the light was applied while the air stream was ON (NB-AL, Air-Light). For the B-condition, values were averaged across Configurations 1 and 6. There was no significant difference between Exc and Inh cells for Resp or zMI, but the RF of Exc cells was larger than that of Inh Cells [*F* (1,4) = 9.85, *p* = .035, η^2^ = .71] (Fig. 4C-E). For the NB-condition, the NB-AL cell population from Condition 4 was compared with cells active for identical Arb events in Configurations 3 and 5 where air was being applied with no light stimulation (NB-A, Air). A two-way RM-ANOVA, with Cell-Type (Exc and Inh) and Event-Type (A and AL) as within-subjects factors, was used. None of the main or interaction effects was significant for any of the variables (Fig. 4F-H).

Cell populations corresponding to the three sensorimotor events B-L, NB-A, and NB-AL were then compared for complementation when taken collectively and pooled across Exc and Inh cell types (Fig. 4J). There were separate populations of cells active around the three events, with NB-A having the largest percentage of complementary cells [RM-ANOVA with Event-Type as a within-subjects factor: *F* (2,8) = 8.99, *p* = .009, η^2^ = .69]. When events were considered two at a time, a three-way RM-ANOVA was conducted, with Event-Type-Pair (e.g., B-L::NB-AL), Population-Type (Comp1, Conj, and Comp2), and Cell-Type (Exc and Inh) as within-subjects factors. There was a significant three-way interaction [*F* (4,16) = 5.83, *p* = .004, η^2^ = .59]. Simple two-way interactions were then examined, and the only difference was between Event-Type-Pair and Population-Type for Inh cells [*F* (4,16) = 16.71, *p* = .005, η^2^ = .81]. Simple simple main effects of Population-Type were then individually determined for all Event-Type-Pairs, and all were significant [B-L::NB-A: *F* (2,8) = 54.13, *p* < .001, η^2^ = .93; B-L::NB-AL: *F* (2,8) = 41.41, *p* < .001, η^2^ = .91; NB-A::NB-AL: *F* (2,8) = 12.12, *p* = .004, η^2^ = .75]. *Post hoc* comparisons revealed that for the first two Event-Type-Pairs (B-L::NB-A and B-L::NB-AL) involving the B and NB conditions there were more complementary Inh cells than conjunctive Inh cells. This was not the case, however, for the third Event-Type-Pair (NB-A: :NB-AL) involving events in only the NB-condition (Fig. 4J). For the Exc cells, the percentage of complementary cells was significantly larger than the percentage of conjunctive cells (Fig. 4J) for all Event-Type-Pairs. These results suggest a segregation of cell populations across the B and NB conditions for both Exc and Inh cells active around light stimuli. Furthermore, there were cells active around light stimuli which were not active around air (Fig. 3I).

### Conjunction and complementation of cell populations characterize stable and dynamic cellular representations of sensorimotor events

The dynamics of the recruitment of cells across different sensorimotor events in all behavioral configurations was analyzed by finding the conjunction and complementation of cell populations active around all sensory and Arb events after pooling across Exc and Inh cell types (Fig. 5A-B). Agglomerative hierarchical clustering of the average heatmaps was then used to identify relationships among conditions (Fig. 5C-E). The clustering analysis showed a segregation of cellular populations across the B and NB conditions, either with clear clustering or branch separation in the dendrograms within the same clusters. However, there was some mixing of populations across the B and NB conditions for the Arb events.

**Figure 5.**
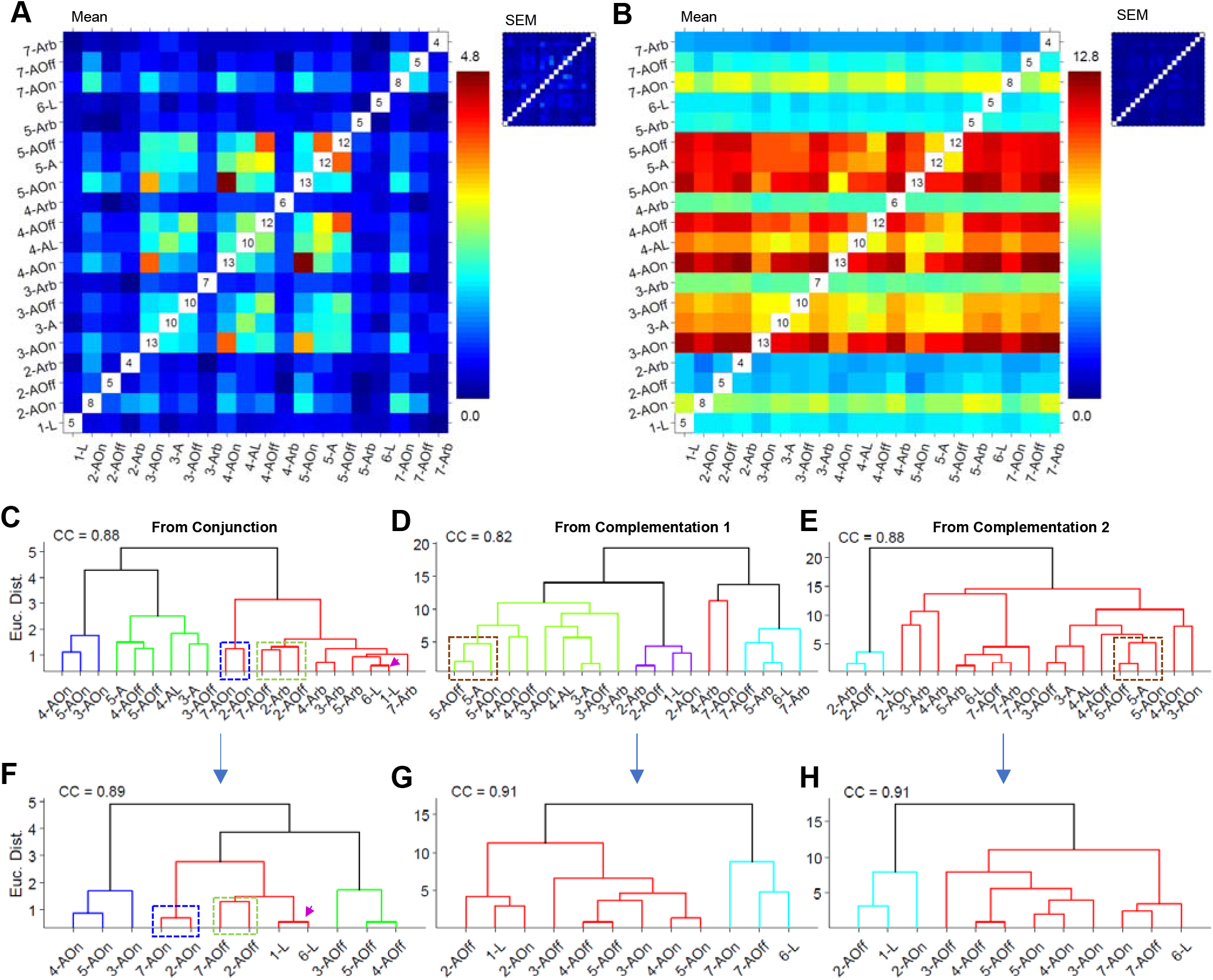
Separate subgroups of cells are active for distinct sensorimotor events. **(A-B)** Average and SEM (insets) heatmaps (over 5 mice) of conjunction and complementation of cell groups active in the Brake and No-Brake conditions around sensory air, light, and Arb events. **(C-E)** Dendrograms resulting from agglomerative hierarchical clustering of the average conjunction and complementation heatmaps/matrices. **(F-H)** Dendrograms resulting from agglomerative hierarchical clustering of the average conjunction and complementation heatmaps/matrices after excluding arbitrary events. CC indicates cophenetic correlation.

The clustering from the conjunction heatmap (Fig. 5C) revealed that distinct subgroups of cells were active for distinct sensorimotor events. For example, the blue cluster shows the closeness of cells active around AOn in the NB-condition (Configurations 3, 4, and 5). A similar relationship existed for the B-condition (Configurations 2 and 7), as can be seen in the dendrogram branch in the red cluster enclosed within the dotted blue line. The green cluster represents cells active around AOff in the NB-condition, and the branch in the red cluster enclosed within the dotted green line represents similar cells for the B-condition. The branch indicated by the magenta arrowhead shows the closeness of populations active around light in the B-condition (Configurations 1 and 6). The cell populations active around Arb events in the NB-condition (3-Arb, 4-Arb, and 5-Arb), however, were closer (in dendrograms) to cell populations active in the B-condition. When the Arb, A, and A-L events were excluded from clustering, a more distinct grouping of cell populations was observed, with clearer separation between distinct sensorimotor events (Fig. 5F) and between the B and the NB conditions.

The clustering of both complementation heatmaps (Fig. 5D-E) grouped together sensorimotor events that either belonged to the same Configuration or that happened close in time. For example, in the purple and cyan clusters in Figures 5D and 5E, respectively, 1-L was grouped with the 2-AOff and 2-Arb events (for the B-condition). Similarly, the green cluster in Figure 5D shows grouping of sensorimotor events in the NB-condition with branch proximity of sensorimotor events belonging to the same Configuration (e.g., 5-AOff, 5-A, 5-AOn, enclosed within the dotted brown line). The cyan cluster in Fig. 5D shows grouping of events in Configurations 6 and 7 with the 5-Arb events from the NB-condition. In both heatmaps for complementation, when Arb, A, and A-L events were excluded, a clear separation between the B and NB conditions was seen. Furthermore, within branches, grouping of sensorimotor events occurred based on the Configuration (or Configurations) that occurred closer in time (Fig. 5G-H). For example, 1-L is grouped with 2-AOn and 2-AOff in Fig. 5G.

The conjunction and complementation of the above-mentioned populations was also studied for cells that responded around voluntary motion onset and offset events (i.e., when a mouse started or stopped moving at a time when no explicit sensory stimuli was applied). In particular, during the no-stimulus intervals of the NB-condition, animals sometimes walked or ran spontaneously. These events were identified and subdivided into components in which an animal started locomoting (motion onset, MOn) or stopped locomoting (motion offset, MOff). For the 5 animals, the number of MOn events detected was 8, 21, 19, 20, and 19, and the number of MOff events was 35, 24, 31, 37, and 39. Cellular activity around these events (1.5 s pre- and post-event) was assessed by generating raster plots and PETHs to identify voluntary motion-related cellular responses. Both Exc and Inh cell responses (Fig. 6A) were identified. Rate vector maps (Fig. 6B, top row) and population correlation plots (Fig. 6B, middle and bottom rows) showed the binary nature of the responses. The effect of Event-Type (MOn vs. MOff) and Cell-Type (Exc vs. Inh) on the percentages of responsive cells was examined with a RM-ANOVA. There was a significant interaction effect [*F* (1,4) = 45.73, *p* = .002, η^2^ = .92], with a greater number of cells showing an Exc than an Inh response to MOn (*p* = .025), and vice versa for MOff (*p* = .002) (Fig. 6C). The same analysis for RF revealed a significant effect of Event-Type (Fig. 6D) [*F* (1,4) = 8.12, *p* = .046, η^2^ = .67]. The fidelity of the cells responding to MOn was significantly greater than that of the cells responding to MOff. There was no significant main effect or interaction between factors for zMI values (Fig. 6E). These results suggest more cells responded during locomotion, and these cells had a more robust response to MOn than MOff.

**Figure 6.**
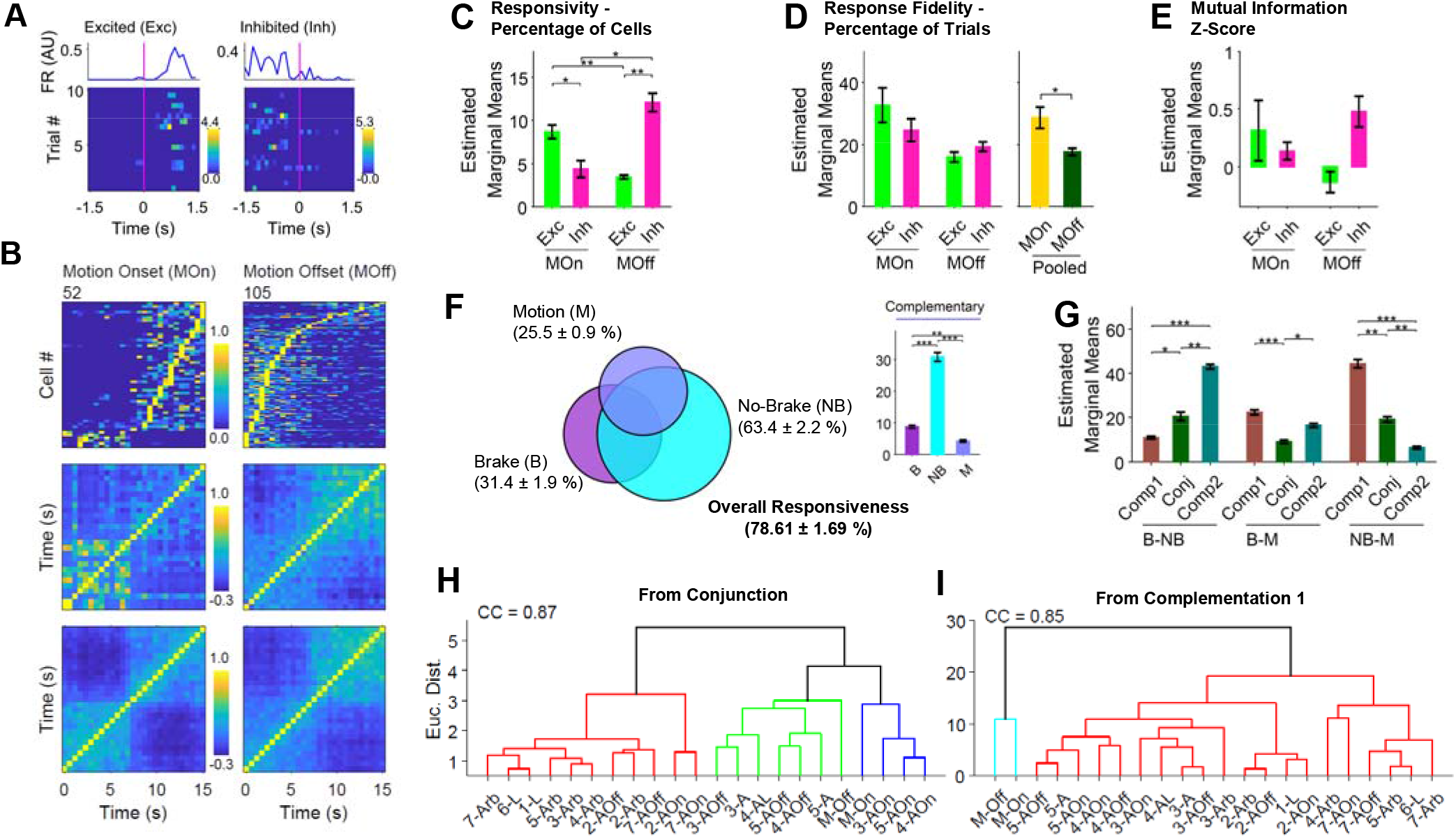
Separate cell groups at movement onset (MOn) and offset (MOff) events compared to other sensorimotor events. **(A)** Raster plots of representative Exc and Inh cells to MOn. The magenta line at t = 0 indicates motion onset. The blue lines show means over trials. **(B)** Rate vector maps (top row) and corresponding population vector correlation plots (middle row) of responsive cells from a representative animal for MOn and MOff events. Average population correlation plots (bottom row) for 5 animals. **(C-E)** Results of two-way RM-ANOVA for assessing responsivity, response fidelity, and mutual information z-score for MOn vs. MOff and Exc vs. Inh cells. **(F)** Overall responsivity, conjunction, and complementation for Brake, No-Brake, and Motion conditions. **(G)** Results of two-way RM-ANOVA for assessing the percent of cells across population types (Comp1, Conj, and Comp2) and condition pairs e.g., Brake and Motion (B-M). **(H-I)** Dendrograms resulting from agglomerative hierarchical clustering of the average conjunction and complementation heatmaps/matrices when MOn and MOff events were included. CC indicates cophenetic correlation. For all statistical tests, *n* = 5 mice. Error bars = SEM. \**p* < .05, \*\**p* < .01.

To examine whether cells exhibiting responses to MOn and MOff (M-condition) responded to events during the B-condition as well as non-voluntary motion events in the NB-condition, cells were pooled across events (AOn, AOff, Arb, L, A, and A-L), response types (Exc vs. Inh), and stimulus types (air and light). Overall, 78.61 ± 1.69 % of cells were responsive, with 8.8 ± 0.4 %, 30.8 ± 1.4 %, and 4.2 ± 0.3 % active in the B-, NB-, and M-conditions, respectively, and 6.9 ± 0.9 % responding during all three conditions (Fig. 6F). When taken collectively, the percentage of complementary cells in the NB-condition was significantly larger than in the B and M conditions (*p* < .001 for both comparisons), and the percentage in the B-condition was larger than in the M-condition (*p* = .003) (Fig. 6F) [RM-ANOVA, *F* (2,8) = 210.72, *p* < .001, η^2^ = .98]. When taken two at a time, the percentages of complementary and conjunctive cells were compared using a two-way RM-ANOVA, with Condition-Pair (e.g., B-M, NB-M) and Population-Type (Conj, Comp1, and Comp2) as within-subjects factors. There was a significant interaction between the two factors [*F* (4,16) = 210.78, *p* < .001, η^2^ = .98]. *Post hoc* comparisons revealed that, for the B-M comparison, the percentage of conjunctive cells was significantly smaller than the percentage of complementary cells (Fig. 6G, *p* = .012 and *p* < .001). However, there was a larger overlap of cells between the NB and M conditions, with a larger percent of complementary cells in the NB-condition (Fig. 6G, NB-M comparison, *p* = .001, .0003, and .0047).

The conjunction and complementation of cells responding to MOn and MOff events was also analyzed with all other sensorimotor events in the B and NB conditions. Cell populations for MOn and MOff were closer to AOn and AOff events, respectively, in the NB-condition, as revealed by clustering of the conjunction heatmap (Fig. 6H). Clustering of the complementation heatmaps showed that MOn and MOff formed a separate cluster, suggesting that the complementary cells related to voluntary motion were distinct compared to other cell populations.

Taken together, these results suggest that there are separate cell populations which are active for distinct sensorimotor events in different behavioral configurations. Where the conjunctive cell populations indicated stable cellular representations, complementary cell populations featured the dynamics of recruitment across different sensorimotor events. This possibility was analyzed next by doing trial-by-trial analyses on the activity of cells.

### Trial-wise turnover of cellular recruitment reveals a larger percentage of complementary cells for the No-Brake condition in early trials

To examine how cells were recruited during individual trials during the presentation of air, light, and arbitrary events in relation to the B and NB conditions, responsive cells (firing rate > 0) were identified for each trial. The percentage of responsive cells in each trial was found for the 20 different sensorimotor events (see Fig. 7A). Across all trials for all events, 99.7 ± 0.1% of cells were responsive. Nevertheless, for any given event (Fig. 7A, magenta line), an average of 17.3 ± 0.1% (range: 13.07-23.02, median: 17.34) of cells were responsive in any given event. To statistically compare the percentage of responsive cells across the 20 events and 10 trials, a two-way RM-ANOVA was used. There was a significant effect of Event-Type on the percentage of responsive cells (Fig. 7C) [*F* (19,76) = 2.10, *p* = .012, η^2^ = .34]. *Post hoc* comparisons revealed differences in means for a few events (Fig. 7C). Consistent with previous observations, a larger percentage of cells responded to sensory stimuli during the NB-condition, and there were abrupt increases in the percentage of responsive cells at Brake to No-Brake transitions, the condition permitting running onset in Configuration 3.

**Figure 7.**
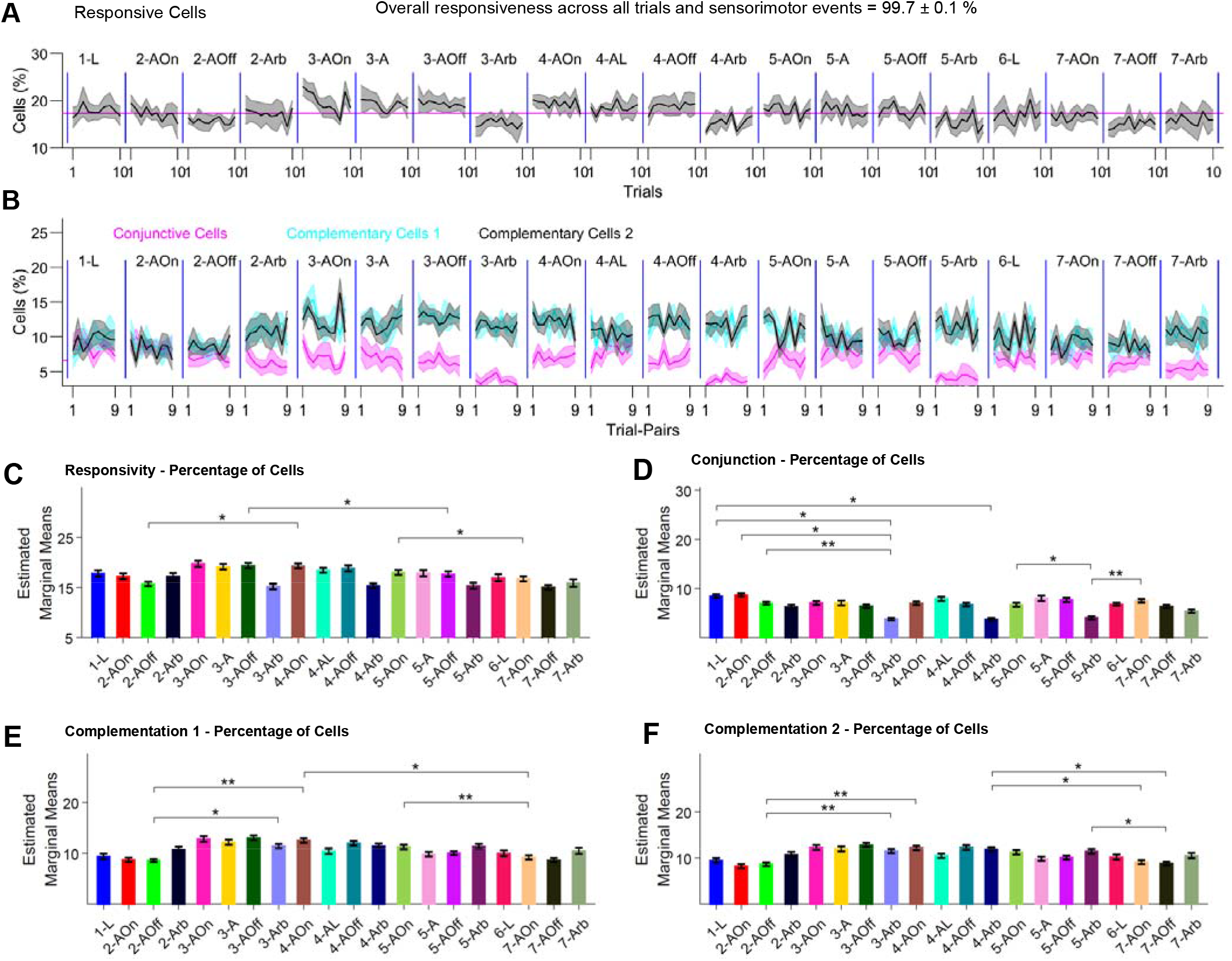
Adjacent trial-wise dynamics of cellular responses and conjunction/complementation in cellular populations. **(A)** Percent of responsive cells (firing rate > 0) over trials for 20 types of cellular responses to different sensorimotor events exhibited across the 7 behavioral configurations. **(B)** Percent of complementary and conjunctive cells for pairs of adjacent trials. Pairs 1-9 correspond to Trials 1-2, 2-3, 3-4, etc. up to 9-10. **(C)** Result of two-way RM-ANOVA assessing responsivity across 20 events and 10 trials. **(D-F)** Results of two-way RM-ANOVA assessing the percent of conjunction and complementation across 20 events and 9 trial-pairs. For all statistical tests, *n* = 5 mice. Error bars = SEM. \**p* < .05, \*\**p* < .01.

The dynamics of the percentage of complementary and conjunctive cells was also assessed by considering adjacent pairs of trials (trials 1 and 2, trials 2 and 3, etc.). For the 9 pairs of trials for each of the 20 events, Conj, Comp1, and Comp2 cells were identified (Fig. 7B). Where Conj cells were active in both trials in a trial pair, Comp1 cells were active in the former trial but became silent in the latter trial, and Comp2 cells started silent but became active in the latter trial.

A two-way RM-ANOVA, with Event-Type and Trial-Pair as within-subjects factors, of the percentage of Conj cells revealed a significant effect of Event-Type [*F* (19,76) = 5.87, *p* < .001, = .59]. *Post hoc* tests revealed significantly smaller percentages of Conj cells during Arb events compared to some of the other events (Fig. 7D, see significant *post hoc* comparisons). A similar analysis of the complementary cells showed that percentages of both Comp1 and Comp2 cells were significantly different across Event-Types [Comp1: *F* (19,76) = 2.81, *p* < 0.001, η^2^ = .41; Comp 1: *F* (19,76) = 2.82, *p* < 0.001, η^2^ = .41]. *Post hoc* comparisons revealed a larger percentage of cells in both complementary populations during the NB-condition compared to the B-condition (Fig. 7E-F, see significant *post hoc* comparisons). For the percentage of Comp2 cells, there was also a significant effect of Trial-Pair [*F* (8,32) = 2.41, *p* = .037, η^2^ = .38]. *Post hoc* comparisons did not show any differences in means across trial-pairs. However, if only air and light stimuli were considered, the percentage of Comp2 cells was significantly different across events [*F* (12,48) = 3.79, *p* = .0005, η^2^ = .49] as well as trial-pairs [*F* (8,32) = 2.76, *p* = .019, η^2^ = .41], with trial-pair 1-2 having a larger percentage of Comp2 cells compared to trial-pair 9-10 (*post hoc* comparison, *p* = .021). These results suggest locomotion contributed to more new cells being recruited immediately following a new event (i.e., a new experience).

The conjunction and complementation of cells for all trial-pairs (200 × 200), corresponding to 20 events with 10 trials per event, were qualitatively assessed with heatmaps. The average heat maps for conjunctive and complementary cells are shown in Fig. 8. The heatmap of conjunctive cells (Fig. 8A) shows that separate subgroups of cells were selectively active during different events. For example, see the checkerboard-like pattern in the middle of the map outlined by the thick brown box, which shows that separate subgroups of cells responded to AOn versus AOff (outlined with red boxes). Furthermore, some of the cells that participated during the first B epoch (Configurations 1 and 2) also did so selectively during the second B epoch (Configurations 6 and 7). The two epochs are outlined with purple boxes. Cells that responded to light during the NB-condition had a high degree of overlap with cells that responded to AOff during the NB-condition. To perform clustering on the conjunction heatmap, the heatmap was binned by averaging across trial-pairs (i.e., all 100 pixels corresponding to 10 × 10 trial-pairs within each big black box bounded by thin lines were averaged). The heatmap thus obtained was then subjected to agglomerative hierarchical clustering. This analysis showed that there was a clear segregation within the dendrograms for the B versus NB conditions (Fig. 8C, branches in the dotted green box vs. other branches). Further segregation can be seen in the separate groupings of the sensory and arbitrary events (blue and purple dotted boxes and cyan cluster). The segregation between the B and NB conditions was clearer when Arb and A events were excluded (Fig. 8F).

**Figure 8.**
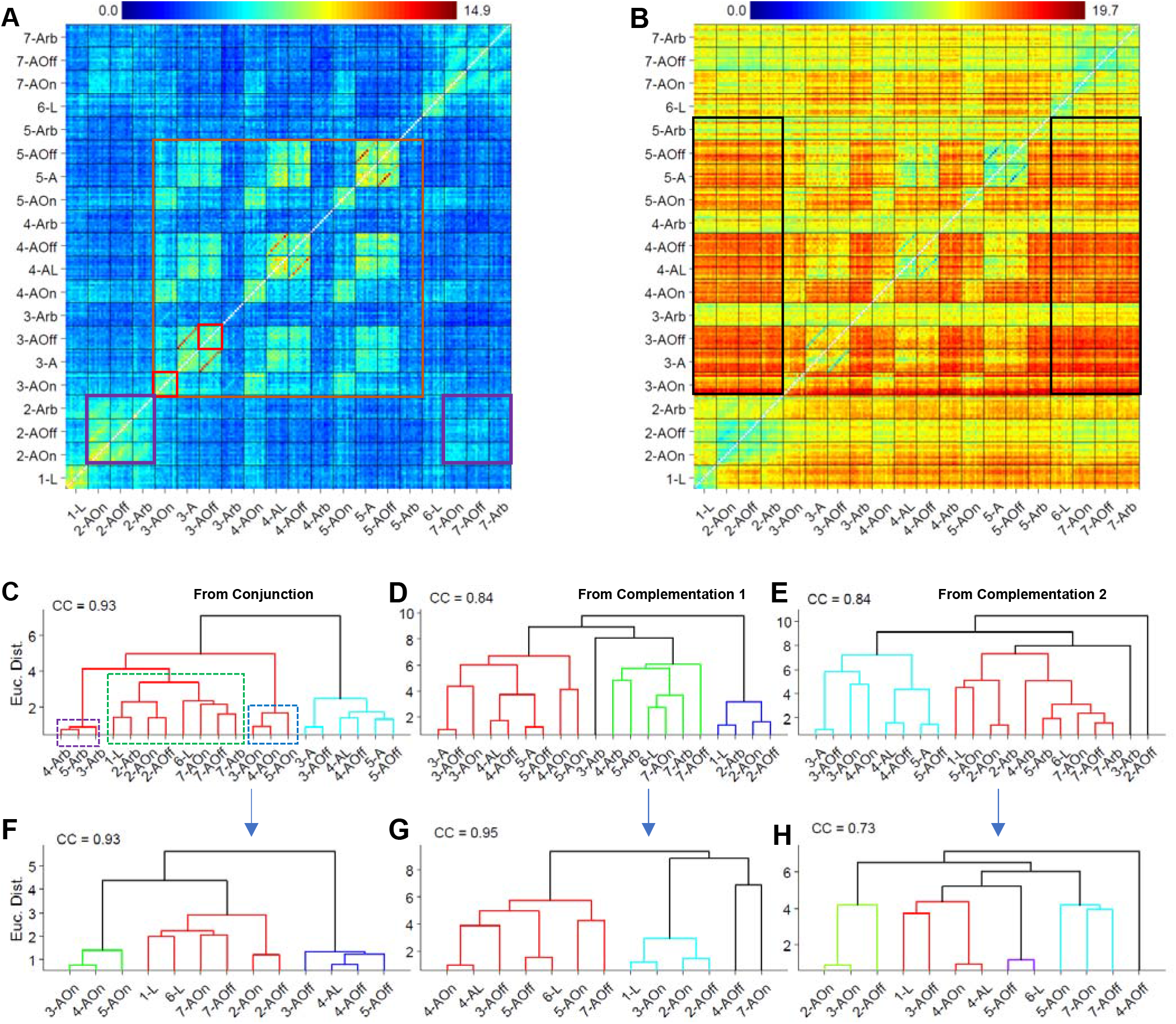
Trial-pair-wise dynamics of conjunction and complementation of cellular populations for all 20 sensorimotor events. **(A)** Heatmap of the average percentage of conjunctive cells (n = 5 mice) between trials from all 20 events. The 200 × 200 pixels (off diagonal) belong to trial-pairs from all 20 events, as shown by the x and y labels. Each big box enclosed by thin black lines shows 10 × 10 trial pairs of the given event. For example, the color value of the pixel in row 1, column 2 indicates the percentage of conjunctive cells for trial 1 and 2 of presentation of the light stimulus. **(B)** Heatmap of the average percentages of complementary cells (n = 5 mice) between trial pairs (like A). Here, for example, the color value of the pixel in row 1, column 2 indicates the percentage of cells present in trial 1 and not present in trial 2. **(C-E)** Dendrograms resulting from agglomerative hierarchical clustering of the average conjunction and complementation heatmaps/matrices. **(F-H)** Dendrograms resulting from agglomerative hierarchical clustering of the average conjunction and complementation heatmaps/matrices after excluding arbitrary events. CC indicates cophenetic correlation.

The heatmap of complementary cells (Fig. 8B) revealed a larger complementation between NB and B conditions (outlined with black boxes). Furthermore, the checkerboard-like pattern in the middle region of the heatmap suggests selective activation during AOn versus AOff during the NB-condition. Clustering of the complementary heatmaps (Comp1 and Comp2) was done in a similar manner as described above, after averaging over trial-pairs and separating the upper and lower triangular regions. Here, the clustering revealed segregation of the B and NB conditions but with some mixing owing to the ongoing recruitment of complementary cells that were similarly active across events or Configurations that happened close in time. The exclusion of Arb and A events before clustering revealed similar results (Fig. 8G-H).

In summary, the results from the trial-by-trial analysis were consistent with findings on the responsiveness of cells averaging over trials. More cells were recruited during the transition from the B to the NB-condition, with overall a greater degree of complementation for the NB-condition. Furthermore, complementation was greater in the initial trials (perhaps due to the novelty of the experience) and became smaller as the experience was repeated. Additionally, distinct smaller groups of cells encoded distinct sensorimotor events.

## Discussion

Calcium imaging documented the activity of hippocampal CA1 cells and revealed different individual and collective cell population responses to sensory stimulation during immobility, locomotion, and the transitions between these behaviors. Head fixed mice stood on a conveyor belt and immobility was encouraged by activating a conveyor belt brake, whereas locomotion could occur when the brake was released. Light and tactile air stimuli were systematically applied during the B and NB conditions. Neuronal excitation or inhibition featured a salt and pepper arrangement suggesting that responsivity was not topographically organized and did not reflect a general function such as immobility versus locomotion or arousal. Overall, ~99% of 2083 cells were active within the imaging session, with ~78% responding in relation to individualized experimental sensory/movement events. On a given trial, combinations of sensory and locomotor events featured subpopulations of active neurons comprising about ~17% of all detected neurons, organized in complementary and conjunctive patterns of activation. From a large population of cells, complementary and conjunctive cells encode relatively novel and familiar experiences, respectively. These results suggest that hippocampal CA1 cells participate in functional networks that distinguish sensory events in relation to ongoing moment-to-moment behavior using a recruitment with replacement strategy.

A central finding of the present study was the individuation of cell responses to events featuring configurations of light, air, locomotion, and immobility. By observing cellular responses to different sensory stimuli during multiple behavioral states in a single paradigm, this study describes the sensorimotor relations of an unprecedented 99% responsive cells. This responsivity could be attributed to the bias of Suite2P software in identifying only active cells. In other words, to the extent that Suite2P failed to identify some cells in the recording frame, particularly cells that were silent across the recording, the 99% figure is an underestimate of the percentage of total CA1 neurons that were responsive. Overall, ~78% of cells (determined from average responses) were tuned to the experimentally determined stimuli and motion events, with a larger percentage of cells active during locomotion versus immobility. Previous studies have reported locomotion-related responses, including sensory-spatial (Herzog et al., 2019; Komorowski et al., 2009; Manns et al., 2007; Moita et al., 2003; Zhao et al., 2020), sensory-temporal (MacDonald et al., 2013; Taxidis et al., 2020), and spatial-temporal (Haimerl et al., 2019; Kraus et al., 2013; Villette et al., 2015) populations with smaller percentages or limited numbers of active cells. For example, (Taxidis et al., 2020) reported sensory-temporal representations in up to 30% active cells, (Alme et al., 2014) reported responses related to 11 different rooms of 30 active cells per animal (freely behaving), and (Grieves et al., 2020) used a 3D environment and described 756 place cells from 13 rats (in contrast to 2083 responsive cells from 5 mice in the present experiment). That the percentage of cells responsive in all conditions in the present study was negligible argues that to fully sample hippocampal cell diversity it is necessary to study the activity of cells during an array of sensorimotor conditions.

Although the percentage (~7%) of conjunctive cells (i.e., cells that were responsive to specific sensory-motor configurations, such that, for a given pair of configurations A and B, C ∈ A and B) was fairly consistent across conditions of immobility and locomotion, the percentages of complementary cells (i.e., cells that were responsive to a specific stimulus or condition, such that C ∈ A not B or C ∈ B not A) were larger during locomotion. Moreover, the complementary cell populations varied with experience (as could be seen when comparing later to earlier trials within a configuration), with the population containing a larger percentage of the total detected cells in the first few trials. This observation supports the idea that ongoing cellular recruitment might depend on experience as well as moment-to-moment behavior, as novel experiences become familiar experiences. The existence of conjunctive populations across similar Configurations (e.g., 2 and 7 OR 3, 4, and 5), additionally argues that familiarity is also encoded in ongoing cell activity.

For both immobility and locomotor related cellular responses, the tactile air stimulus was more potent in eliciting sensory responses than was the visual light stimulus. There were also more cells, with greater response fidelity and stability, that responded to air compared to the light stimulus. This is consistent with a finding in rabbits that repetitive tactile stimulation is more potent than repetitive visual stimulation at modulating hippocampal field potentials (Whishaw and Dyck, 1984). Here, cell responsivity could have been influenced by many factors, including the shorter duration of the light stimulus (200 ms) compared to the air stimulus (5 s); differences in behavioral responses of “freezing” to the stimuli; and differences in autonomic responses to the stimuli, including respiratory and pupillary response changes or slight body movements by the animals. Future experiments using videography combined with calcium imaging can address these possibilities. For locomotion related cell responses, we cannot rule out higher order responses to sensory stimulation or behavior. For example, cellular responses to an AOff event might have some contributions from a previous air or light onset event or vice versa. It should be noted that, although we did not record behavior-concurrent field potentials, a substantial literature reports that electrophysiological signatures change as animals transition between immobility and movement (see introduction).

The cell/behavior relationship described in the present study provides insights into the sensorimotor functions of the hippocampus as a neural network. First, almost no cells were consistently active in association with all sensorimotor events, as might be expected in a system related to general arousal (Green and Arduini, 1954) or reflecting only ongoing movement (Vanderwolf, 1969). Second, cell responses were not topographically organized as is found in primary sensory and motor regions of the neocortex, a finding consistent with the idea that the hippocampus is a network system (Buzsaki, 1996). Third, cells obtain specificity with respect to sensorimotor behaviors by featuring complementary and conjunctive subpopulations. For example, a subpopulation of cells might respond in relation to a tactile air stimulus when a mouse is relatively immobile, whereas a different, complementary subpopulation might respond to the same stimulus when the mouse is moving, and a smaller, conjunctive population might respond to the stimulus in both conditions. Fourth, complementary and conjunctive subpopulation cell numbers appeared relatively fixed around average numbers of 10% and 7%, respectively.

In summary, the complementary/conjunctive specificity of CA1 cells suggests that the hippocampus is a network that encodes ongoing sensorimotor events related to concurrent movement. Such cell-behavior relations are congruent with reports of the relationship between hippocampal field potentials and behavior. Given this complementary and conjunctive cell subpopulation organization, it is not surprising that subpopulations of cells can be secondarily related to features of spatial behavior (O’Keefe and Dostrovsky, 1971; O’Keefe and Nadel, 1978), episodic learning and memory (Eichenbaum, 2004; Eichenbaum and Cohen, 2014), context representation (Bulkin et al., 2020; Rudy, 2009; Smith and Bulkin, 2014), and scene construction (McCormick et al., 2021), as the hippocampus is downstream from the many cortical and subcortical systems that are also involved in these processes.

## Methods

### Animals and surgery

Animal procedures were performed in compliance with protocols approved by the Animal Welfare Committee of the University of Lethbridge and followed the guidelines for the ethical use of animals provided by the Canadian Council on Animal Care. In total, 5 Thy1-GCaMP6s [C57BL/6J-Tg(Thy1-GCaMP6s)GP4.3Dkim/J] (JAX stock #024275) hemizygous transgenic mice were used (3 male, 2 female). This mouse line exhibits stable and homogenous expression of GCaMP6s over a large population of cortical and subcortical excitatory neurons (Chen et al., 2013b; Dana et al., 2014). After cranial window surgery mice were housed in individually ventilated Optimice cages (Animal Care Systems) in temperature-controlled rooms (22 °C) maintained on a 12:12 light/dark cycle (lights on during the day). Food and water were provided *ad libitum*.

Mice were implanted with chronic cranial windows at ~5-7 months of age. Mice were injected subcutaneously with buprenorphine (0.075 mg/kg) 0.5 h before surgery. Immediately prior to surgery, animals were also injected subcutaneously with 0.5 ml of a 5% dextrose, 0.9% saline solution mixed with atropine (3 μg/ml). Animals were anesthetized with isoflurane (2-3% for induction, 1-1.5% for anesthesia) and their body temperature was maintained at 37 °C using a homoeothermic monitoring system and heating pad. Lidocaine (10 mg/kg) was injected under the scalp as a local anesthetic, and a section of skin was removed with scissors to expose the skull. A craniotomy, 3 mm in diameter, was made above the right hippocampus (center at −2.0 mm AP, 1.8 mm ML, relative to bregma). The dura underlying the craniotomy was resected, and a small region of neocortex was removed by aspiration, exposing the underlying white matter. A metal cylinder (3 mm outer diameter, 1.5 mm long) with a glass coverslip attached to the bottom end using optical adhesive (NOA71, Norland) was inserted into the cavity. The coverslip was gently pushed against the tissue to reduce motion, and the cylinder was attached to the skull using cyanoacrylate tissue adhesive (Vetbond, 3M) and dental acrylic. To allow for head-fixation of the mouse, a titanium head-plate was attached to the skull using adhesive cement (C&B Metabond, Parkell). Two rubber rings were attached to the top of the head-plate, forming a well that could be filled with distilled water for imaging with a water-immersion objective. Post-surgery, subcutaneous injections of meloxicam (7.5 mg/kg) and enrofloxacin (10 mg/kg) were administered daily for 2-3 days. Animals were allowed to recover for 3-8 weeks before behavioral training was started. The age of the animals at the time of imaging was ~8 months.

### Animal training and behavioral experiment

Animals were habituated to head-fixation over 3 days. Mice were head-fixed twice per day, with sessions gradually increasing in length from 5 to 30 min. Following habituation, animals were trained to run on a linear conveyor belt 150cm in length while head fixed. The conveyor belt has been previously described (Mao et al., 2017; Mao et al., 2018). The conveyor belt was not motorized; all movement of the belt were generated by the mouse. The belt was made from Velcro material (Country Brook). Training was done in the dark, on a blank conveyor belt devoid of sensory cues. Belts were washed after every training session. Mice were trained for 20-30 min each day to run in response to a continuous, mild air stream applied to the back. A solenoid valve based automated system was used to apply a continuous mild air stream from the building’s compressed air supply via a regulator. To escape this stimulus, the animal was required to run a fixed distance on the belt, after which the air puff ceased. Following an inter-trial interval of 15 s, the air stream would turn on again, initiating the next training trial. The distance the mouse was required to run was increased gradually across training trials, starting each day with 30 cm, and increasing to a maximum of 150 cm (i.e., one full turn of the belt). On a given training trial, if the mouse ran the required distance with an average speed greater than 7 cm/s, the required distance on the next trial was incremented by 7-15 cm. If the mouse failed to achieve the target speed (e.g., due to running too slowly, delayed starting, or repeated starting and stopping), the required distance did not change between trials. Mice were trained until they reached the maximum distance of 150 cm and achieve the target speed on >75% of trials across a training session. All mice reached criterion in 3 training sessions.

After training, mice were tested (re-trained) 10-18 days later and the following day they experienced a 7-configuration sensorimotor behavioral paradigm while activity of hippocampal CA1 neurons was recorded by two-photon calcium imaging. Behavioral data were simultaneously acquired and synchronized with the imaging data using a data acquisition system (Axon Digidata 1322A). These data included the distance traveled on the belt (measured by rotation encoders attached to the shafts of the treadmill wheels). A fixed reference point on the belt was also tracked, using reflective tape attached to the underside of the belt, which activated a photosensor once per belt lap. Air puff onset and offset, as well as brake and visual cue onset and offset, were controlled by a microprocessor board (Arduino Mega). The results of air training and analyses of distance and/or duration tuning from air onset and offset in Configurations 3, 4, and 5 as well as additional analyses will be presented elsewhere.

### Two-photon calcium imaging

A multiphoton microscopy system (Bergamo II, Thorlabs) was used for two-photon calcium imaging. A Ti:Sapphire excitation laser (Coherent) was tuned to 920 nm, with a power of ~100 mW at the sample. Laser scanning was controlled by galvo and resonant scanners. A GaAsP photomultiplier tube was used to detect the green fluorescence signal from GCaMP6s. Samples were imaged through a 16×/0.8 NA water-immersion objective (Nikon). A black fabric cover was wrapped around the objective and head-plate to block contaminating light. For two mice, images of hippocampal CA1 pyramidal neurons were captured from two planes, with the dorsal plane ~125 μm beneath the tissue surface and the second plane 50 μm deeper. Images were gathered at a frame rate of 29.16 Hz, but the effective frame rate for each individual plane was 9.72 Hz (29.16 ÷ 3, because 1 extra fly-back frame was also captured). For three mice images were gathered from only one plane with a frame rate of 29.16Hz. A field of view of 418 × 418 μm was captured with a resolution of 512 × 512 pixels.

### Data analysis

Suite2P (Pachitariu et al., 2017) (http://mouseland.github.io/suite2p) was used to register images and to automatically detect regions of interest (ROIs) assumed to represent individual cell bodies. Raw calcium traces were determined for each ROI by averaging the fluorescence signal from all pixels within the ROI and correcting for neuropil contamination. For each raw calcium fluorescence trace (F), baseline (F_o_) was estimated, and baseline subtracted signal (ΔF/F_o_) was then determined as (F-F_o_)/F_o_. From the ΔF/F_o_ calcium traces, deconvolved spike rates were determined using a first order auto-regressive model and constrained nonnegative matrix factorization (Friedrich et al., 2017). Further analysis of the deconvolved traces was conducted using custom scripts written in MATLAB (MathWorks).

To make the raster plots, a time bin width of 0.11s was used (36 bins for 4 s) and the average firing rate was calculated for each time bin. Response fidelity (RF) of a neuron for a raster was defined as the percentage of trials in which the firing rate was greater than zero for any time bin. Mutual information (MI) between firing rate and time was calculated for each raster using firing rate values from all trials by binning them into four non-overlapping quantiles(Souza et al., 2018) using the following relation

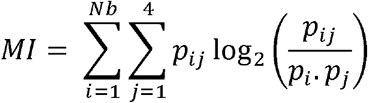

where *p_i_*, and *p_j_* are the probabilities of time bin *i* and firing rate bin *j*, respectively; *p_ij_* is the joint probability between time bin *i* and firing rate bin *j*; *Nb* is the number of time bins i.e., 36.

Peri-event time histogram (PETH) was obtained by averaging the response in the rasters over trials and was used to determine responsiveness of a cell using a two-tailed, two-sample, t-test comparing pre- and post-event firing rates (alpha value 0.05). Rate vector maps were obtained by plotting PETH of all cells after normalizing with the peak of response. The population vector correlation was obtained by finding the Pearson correlation between all pairs of the columns of the rate vector map i.e., each pixel shows correlation between two columns of rate vector map and hence the matrix is symmetric across the diagonal.

### Agglomerative Hierarchical Clustering

This analysis was performed on the conjunction or complementation matrices (heatmaps) when multiple cell groups were considered. Complementation matrices were split into two heatmaps symmetric across the diagonal by using the upper and lower triangular matrices corresponding to Comp1 and Comp2 types of cells before clustering. Based on the conjunction and complementation percentages of each cell group with every other cell group, this analysis found sensorimotor events which were like each other i.e., close to each other with respect to their conjunction/complementation relationships with all other cell groups. The analysis was done by first calculating Euclidian distance for each pair of cell groups considering two rows of a matrix at a time (using Matlab function “pdist”). For example, if a matrix was n × n, there would be 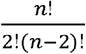 pairs which can then be arranged into a symmetric n × n matrix usually called the dissimilarity matrix owing to smaller Euclidian distance indicating smaller dissimilarity. Matlab function “linkage” was then used to operate on the dissimilarity matrix and define clusters based on the unweighted average distance between clusters. Clusters were then separated (color coded in figures using the Matlab function “dendrogram”) by using a threshold of 70% of the maximum linkage distance between any two clusters. The quality of clustering was assessed by determining cophenetic correlation (CC) reported with each dendrogram in figures.

### Statistical Analysis

Statistical analysis was done in MATLAB® 2018b (Ver 9.5.0.944444) and confirmed for accuracy with IBM® SPSS® (Ver 28.0.1.1). All means in the text are reported ± the standard error of the mean (SEM). Unless otherwise specified, a repeated measures analysis of variance (RM-ANOVA) test was used for statistically comparing differences in measured parameters from *n* = 5 animals. The Shapiro-Wilk test was used to test normality and was done using a file “swtest.m” downloaded from Matlab File Exchange Server (BenSaïda, 2022). The accuracy of the code was confirmed by comparing its output with similar results from SPSS. Minor violations of the normality test and the presence of outliers were ignored (i.e., if some variables did not pass the normality test or had outliers, RM-ANOVA was still performed). The test for sphericity was conducted using Mauchly’s test. Greenhouse-Geisser corrected *p*-values are reported where sphericity was violated. All *post hoc* pairwise comparisons were conducted either using Tukey’s Honestly Significant Difference (HSD) test or the Bonferroni correction with an alpha = .05.

For multi-way RM-ANOVA tests, an alpha level of .05 was used to determine significance at the first level of analysis. For the second level of analysis, *p*-values were adjusted using the Bonferroni correction (e.g., if there was a significant three-way interaction, simple two-way interactions were examined at the second level of analysis, and alpha was divided by the number of two-way interactions tested). Similarly, at the next level of analysis (if required), alpha was further divided by the number of statistical comparisons made (e.g., if there were three simple simple main effects to be tested at the third level of analysis, an alpha level of 0.025/3 = 0.0083 was used). *, **, and *** indicate *p*-values less than .05, .01, and .001, respectively. Because of the small sample size (*n* = 5 animals), effect size is reported along with *p*-values as partial η^2^ = (Sum of Squares _effect_)/(Sum of Squares _effect_ + Sum of Squares _error_).

## Acknowledgements

We thank Dr. JianJun Sun for performing animal surgeries, Di Shao and the University of Lethbridge Animal Care Services staff for animal husbandry, and HaoRan Chang and Drs. Bruce McNaughton, Javad Karimi, Hardeep Ryait, and Žaneta Navrátilová for useful discussions regarding the preparation of this manuscript. We thank Adam Neumann and Dr. Maurice Needham for technical assistance with two-photon microscopy. We also thank Dr. Bryan Souza for providing MATLAB code for calculating mutual information.

## Author Contributions

All authors participated in the design of this study, with S.I. and I.Q.W. as the main contributors. S.I. and B.B.M. performed the experiments. S.I. performed the data analyses with participation of the other authors. S.I. and I.Q.W. wrote the manuscript, which all authors commented on and edited. M.H.M. supervised the study.

